# Multi-attribute characterisation of mRNA via MazF endoribonuclease and LC-MS workflows

**DOI:** 10.64898/2026.09.23.753728

**Authors:** Emma. N. Welbourne, Gareth R. Owen, Lauren P. Wright, Sara Trabulo, Zoltán Kis, Mark J. Dickman

**Affiliations:** School of Chemical, Materials and Biological Engineering, University of Sheffield, Sheffield, UK; Analytical Sciences, BioPharmaceuticals Development, BioPharmaceuticals R&D, AstraZeneca, Cambridge Biomedical Campus, Cambridge CB2 0AA, United Kingdom; Department of Chemical Engineering, Imperial College London, London, UK

**Keywords:** mRNA vaccines/therapeutics, mass spectrometry, critical quality attributes, mRNA, MazF

## Abstract

The rapid expansion of mRNA-based medicines has driven demand for robust, high-resolution analytical methods capable of characterising critical quality attributes including sequence identity, 5’ capping efficiency, and poly(A) tail length and heterogeneity. Here, we present two complementary liquid chromatography-mass spectrometry based workflows for mRNA characterisation, both based on site-specific digestion with the endoribonuclease MazF from *Escherichia coli*. A “middle-up” approach, optimised via specific “5’-ACA” cleavage enables rapid characterisation via oligoribonucleotide mass mapping providing complete sequence coverage and simultaneous assessment of 5-capping efficiency and 3’-poly(A) tail length and heterogeneity. This workflow was successfully applied to three different mRNA constructs, NLuc, eGFP and FLuc mRNAs, achieving sequence coverages of 100%, 87%, and 74%, respectively. A complementary “bottom-up” approach, employing less specific “5’-ACX” cleavage enables tandem mass spectrometry-based sequencing of shorter oligoribonucleotides and detailed characterisation via mRNA sequence mapping. Applied to SARS-CoV-2 Spike Protein mRNA, this method yielded 50% sequence coverage based on unique oligoribonucleotides with MazF alone, extended to 90% by combining MazF, partial RNase T1, and partial RNase U2 digests. Together, these workflows provide a flexible, orthogonal platform for both high-throughput quality assessment and in-depth primary sequence characterisation of mRNA vaccines and therapeutics.

## Introduction

mRNA has recently been demonstrated as highly efficacious in the development of a new class of medicines, through the development and global approval of the Comirnaty (Pfizer/BioNTech) and Spikevax (Moderna) vaccines against SARS-CoV-2.^1,2^ Furthermore, mRNA based-medicines have the potential for a wide range of treatments, beyond vaccines and infectious diseases, in areas such as cancer, metabolic disorders, cardiovascular conditions and autoimmune diseases.^3–5^

mRNA drug substance is produced via *in vitro* transcription, generally by using bacteriophage T7 RNA polymerase. Analytical methods are key to the development of mRNA-based vaccines and therapeutics, as well as underpinning manufacturing through assessment of batch-to-batch reproducibility and quality control (QC) testing for drug release. There is therefore currently significant demand for the development and implementation of improved analytical methods for mRNA that can provide high level detail of construct characteristics and impurity identification, as well as well-optimised alternatives for more rapid, high-throughput QC checks.

mRNA drug substance contains five key structural components that are essential to its stability and efficacy: a 5’ cap, a 5’ untranslated region, a coding sequence, a 3’ untranslated region and a 3’ poly(A) tail.^6–9^ Therefore, these components are listed as critical quality attributes (CQAs) of mRNA drug substance: mRNA identity, 5’ capping efficiency and poly(A) tail length and heterogeneity.^7,10–13^ Whilst conventional Sanger or next-generation sequencing methods are still employed in the analysis of these CQAs, liquid chromatography interfaced with mass spectrometry (LC-MS) has emerged as a powerful, orthogonal tool. These LC-MS-based methods are direct, providing an unbiased and accurate evaluation of the mRNA primary sequence and its modifications without the need for conversion to cDNA or amplification, whilst also providing high-throughput sequence identification and sensitive impurity detection.

Recently, a number of workflows have been developed based on the use of site-specific endonucleases, paired with powerful mass spectrometry-based analysis for the rapid characterisation of large mRNA constructs. The majority of these employ “bottom-up” approaches, whereby the large mRNA is digested into smaller oligoribonucleotides prior to LC-MS and tandem mass spectrometry (MS/MS) analysis.^14–19^ This type of workflow is performed using instruments with the capability of accurate mass analysis, whilst the MS/MS mode is used to effectively sequence and identify oligoribonucleotides. Although the analysis of capping, poly(A) tails and mRNA sequence predominantly use these bottom-up approaches, there have been some studies that have employed intact mass workflows for the analysis of mRNA purity. In particular, characterisation of the length and heterogeneity of poly(A) tails via intact mass analysis has been successful.^20,21^

To achieve efficient and effective sequence mapping via LC-MS/MS, it is necessary for the mRNA of interest to be cleaved in a way such that the resulting oligoribonucleotides are of both the ideal length for LC-MS/MS analysis (<50 nt) and are unique sequences that map only once to the mRNA sequence. This poses a challenge for designing direct sequencing workflows, which has been addressed through the use of parallel RNase digests, partial RNase digests and the use of lower frequency enzymes.^14–16,19^

One lower frequency enzyme that has already been employed in an mRNA mapping workflow is the sequence-specific endoribonuclease MazF, which forms the toxin component of an *E. coli* toxin-antitoxin system,^22^ and is reported to cleave at 5’-ACA motifs^14,23,24^ or 5’ or 5-NAC where “N” is preferentially U or A.^22^ In this work, we present two complementary LC-MS-based methods that exploit the site-specific cleavage of mRNA using *E. coli* MazF. We first optimise digest conditions to create a “middle-up” workflow, via specific 5’-ACA cleavage of mRNA and intact mass analysis of the corresponding oligoribonucleotide fragments. We deploy this as a rapid quality assessment method for evaluating mRNA identity, including assessment of the poly(A) tail and quantification of 5’ capping efficiency. Secondly, a more conventional “bottom-up” workflow was developed for mRNA sequence mapping. We optimise conditions for cleavage with lower specificity at 5’-ACX sites (where “X” can be any nucleobase) and in conjunction with MS/MS analysis. This is employed in high confidence mRNA sequence mapping and can additionally provide insight into ribonuclease specificity and potential secondary/tertiary mRNA structure.

## Experimental

### Chemicals

Water (UHPLC MS grade, Thermo Scientific), acetonitrile (ACN, UHPLC MS grade, Thermo Scientific), 1,1,1,3,3,3-hexafluoro-2propanol (HFIP, >99.8% Fluka LC MS grade), dibutylamine (DBA, ADD), *E. coli* interferase MazF enzyme and 5x buffer (Takara), NLuc mRNA with CleanCap AG (Aldevron).

### Synthesis and purification of mRNA

mRNA synthesis via *in vitro* transcription (IVT) was performed using linearized plasmid DNA (GenScript) at 2×10^−5^ mM, coding for CSP, eGFP or FLuc. This was combined with DNA-dependent RNA polymerase of T7 bacteriophage (Roche) and ATP, CTP, GTP and UTP (Roche) in an equimolar ratio at 10 mM concentration. The reaction was further supplemented with the standard reaction buffer recommended by the enzyme manufacturer. Inorganic pyrophosphatase (Roche) at 2.9×10^−3^ mM was added to the reaction mixture to prevent magnesium pyrophosphate precipitation. RNase inhibitor (Roche) was added at 2.1×10^−4^ mM to maintain an RNase-free environment in the reaction mixture. The reaction was incubated at 37 °C for 2 hours. Following IVT, template DNA was removed by the addition of DNase I and RNA was purified using silica columns as previously described.^25^ RNA concentrations were determined using a NanoDrop^TM^ 2000c spectrophotometer (ThermoFisher Scientific) by absorbance at 260 nm normalized to a path of 1.0 cm.

### MazF digestion of mRNA

MazF digests were performed under the conditions of 10 U enzyme:1 µg of mRNA. For -ACA-directed cleavage of mRNA a final concentration of 2x MazF buffer and an incubation time of 4 hours at 37 °C was used. For less specific, -ACX cleavage of mRNA a final concentration of 1x MazF buffer and varying ratios of MazF enzyme:1 µg of mRNA were used with an overnight incubation at 37 °C.

### LC-MS analysis of mRNA digests

mRNA digest samples were analysed by ion-pair reversed-phase HPLC in conjunction with mass spectrometry. A Vanquish binary gradient UHPLC system (Thermo Fisher Scientific), using a DNAPac RP column (2.1 mm I.D., Thermo Fisher Scientific), was implemented for chromatography. Chromatograms were generated using UV detection at a wavelength of 260 nm. Chromatographically separated mRNA digests were interfaced with an Orbitrap Exploris 240 MS instrument (ThermoFisher Scientific) for subsequent intact mass or MS/MS analysis.

The chromatographic analysis of MazF digests was performed using the following conditions: buffer A 10 mM DBA and 50 mM HFIP; buffer B 10 mM DBA, 50 mM HFIP, and 50% ACN. A flow rate of 0.25 mL/min was used.

For the middle-up workflow the gradient started at 30% buffer B, followed by linear extension to 35% buffer B over 1 minute. Buffer B was then increased to 50% over 9 minutes (curve 3). A temperature of 50 °C was used. MS data was acquired in “Profile” mode using the “Intact Protein” application mode and the “Low Pressure” mode. Spectra were captured at Orbitrap resolutions of 15,000 and 120,000 with a scan range of 450–2500 m/z, RF lens 75%, a normalised AGC target of 100%, and a 100 ms maximum injection time. 3 microscans were performed in negative mode.

For the bottom-up workflow the gradient started at 5% buffer B and linearly increased to 10% buffer B over one minute, followed by a linear extension to 40% buffer B over 29 minutes. A temperature of 80 °C was used. MS/MS data was collected using data dependent acquisition in full scan negative mode with an MS1 resolution of 120,000 and a normalised AGC target of 200%. MS1 ions were selected for higher energy collisional dissociation. MS2 resolution was set at 30,000 with an AGC target of 100%, isolation window of 3 m/z, scan range of 150–2000 m/z and normalised stepped collision energies 17, 20 and 23%.

### LC-MS data analysis

LC-MS data was analysed using BioPharma Finder v.5 (Thermo Fisher Scientific). using the “Intact Mass Analysis” experiment type. Isotopically unresolved LC-MS data acquired at 15,000 resolution was processed using the ReSpect™ deconvolution algorithm, whereas the isotopically resolved 120,000 resolution data was analysed using the Xtract™ deconvolution algorithm. Mass deconvolution employed the “Sliding Windows” “Source Spectra” method. Sliding window parameters were adjusted to accommodate the chromatographic peaks. The “Peak Model” was set to “Nucleotide” with a “Negative Charge” specified for deconvolution. Digest fragments were identified against predicted 0 and 1 missed cleavage species from *in silico* digestion of the mRNA sequence (<2 Da from average mass deconvolution). 5’ cap analysis was performed using the Xtract™ deconvolution across the chromatographic peaks containing the 5’ terminal MazF -ACA cleavage fragments. Capping efficiency was calculated in a label-free approach from the sum intensity of the 5’ terminal MazF oligonucleotide fragment containing a Cap-1 structure as a percentage of the sum of the intensities of the Cap-1 and triphosphate-containing oligonucleotides. Extracted ion chromatograms for the Cap-1 and triphosphate containing species were generated in FreeStyle v.1.8 (Thermo Fisher Scientific) using the most abundant charge state for each species. Poly(A) tail analysis was performed using the ReSpect™ deconvolution algorithm across the chromatographic peak containing the 3’ terminal MazF fragments. Tail lengths were determined based on the calculated 3’ terminal ACA cleavage fragments, differing in the number of As in the poly(A) tract.

### LC-MS/MS data analysis

Data analysis was performed in BioPharma Finder v.5.2 (BPF, Thermo Fisher Scientific), using the “Oligonucleotide Analysis” module. For MazF digests an “MS Noise Level” of 10,000 and an “S/N Threshold” of 50 was selected for component detection. To identify large fragment ions, the maximum oligonucleotide mass was set to 150,000 Da, minimum confidence at 0.5 and mass accuracy at 10 ppm. The ribonuclease selection was set to “mazF”, with “Custom Specificity” selected and “-ACA,-ACC,-ACG,-ACU,-AUA” inputted. The specificity level was set at “high”. The phosphate location was set at “none”, so 3’ OH termini was the default.

For RNase T1 and U2 digests “Enable Automatic Parameter Values” was selected for component detection. The maximum oligonucleotide mass was set to 30,000 Da, minimum confidence at 0.5 and mass accuracy at 10 ppm. The ribonuclease selection was set to RNase T1/RNase U2, default specificity (G-) was used for RNase T1 and “Custom Specificity” selected for RNase U2 with “A-,G-“ inputted. The specificity level was set at “high”. The phosphate location was set at “none”, so 3’ OH termini was the default.

For all constructs, phosphorylation and cyclic phosphorylation were set as variable modifications of the 3’ terminal in the sequence manager containing the RNA sequence (excluding the poly(A) tail). Random RNA sequences of the same length and GC content were included in the sequence manager in addition to the correct RNA sequence, these were generated by the random sequence generator in the BPF sequence editor. For data processing and review, additional filters were included: “Identification” = “does not contain nonspecific”, “does not contain nonunique”; “Mod” = “does not contain None”; “Nonunique Seq” = “≤1”; “Δ ppm” = “≤20”, “≥−20”;””Conf. Score” = “≤90”; “Best ASR” = “≥2.0”; “ID Type” = “contains MS2”; “Mono Mass Exp.” = “>0”. All oligonucleotide identifications from BPF are shown in the Supporting Information (Supporting Table S1). Sequence maps produced by BPF were further processed using our in-house software tools to produce linear and spiral maps.^19^

### Cleavage site analysis

Data analysis was performed in BPF (Thermo Fisher Scientific), using the “Oligonucleotide Analysis” module with “Enable Automatic Parameter Values” checked. The maximum oligonucleotide mass was set to 150,000 Da, minimum confidence at 0.5 and mass accuracy at 10 ppm. The ribonuclease selection was set to “Nonspecific”. The specificity level was set at “high”. The phosphate location was set at “none”, so 3’ OH termini was the default.

Phosphorylation and cyclic phosphorylation were set as variable modifications of the 3’ terminal in the sequence manager containing the RNA sequence (excluding the poly(A) tail). For data processing and review, additional filters were included: “Mod” = “does not contain None”; “Δ ppm” = “≤20”, “≥−20”;”“Conf. Score” = “≤90”; “Best ASR” = “≥2.0”; “ID Type” = “contains MS2”; “Mono Mass Exp.” = “>0”.

Cleavage specificity analysis was performed using a custom Python-based workflow. Oligonucleotide identifications from the nonspecific cleavage search were mapped onto the theoretical mRNA sequence following extraction of cleavage start coordinates from reported positional ranges. For each identification, the nucleotide identities at positions −5 to −1 upstream and +1 to +5 downstream of the inferred cleavage site were extracted and appended to the identification table. Position frequency and probability matrices were generated (unweighted and MS-area-weighted). Cleavage site sequence preferences were visualised using sequence logos generated from the position probability matrix.

## Results and Discussion

### MazF-based workflows for mRNA characterisation

In this study we develop two alternative LC-MS workflows for characterising mRNA CQAs: a middle-up, rapid method for multi-attribute analysis and a bottom-up method for more detailed characterisation and identity analysis. Both of these workflows utilise mRNA digestion via *E. coli* MazF, followed by oligoribonucleotide separation by ion-pair reversed-phase HPLC (IP-RP-HPLC) in conjunction with mass spectrometry analysis. A schematic overview of the two workflows is shown in Figure 1.

**Figure 1:**
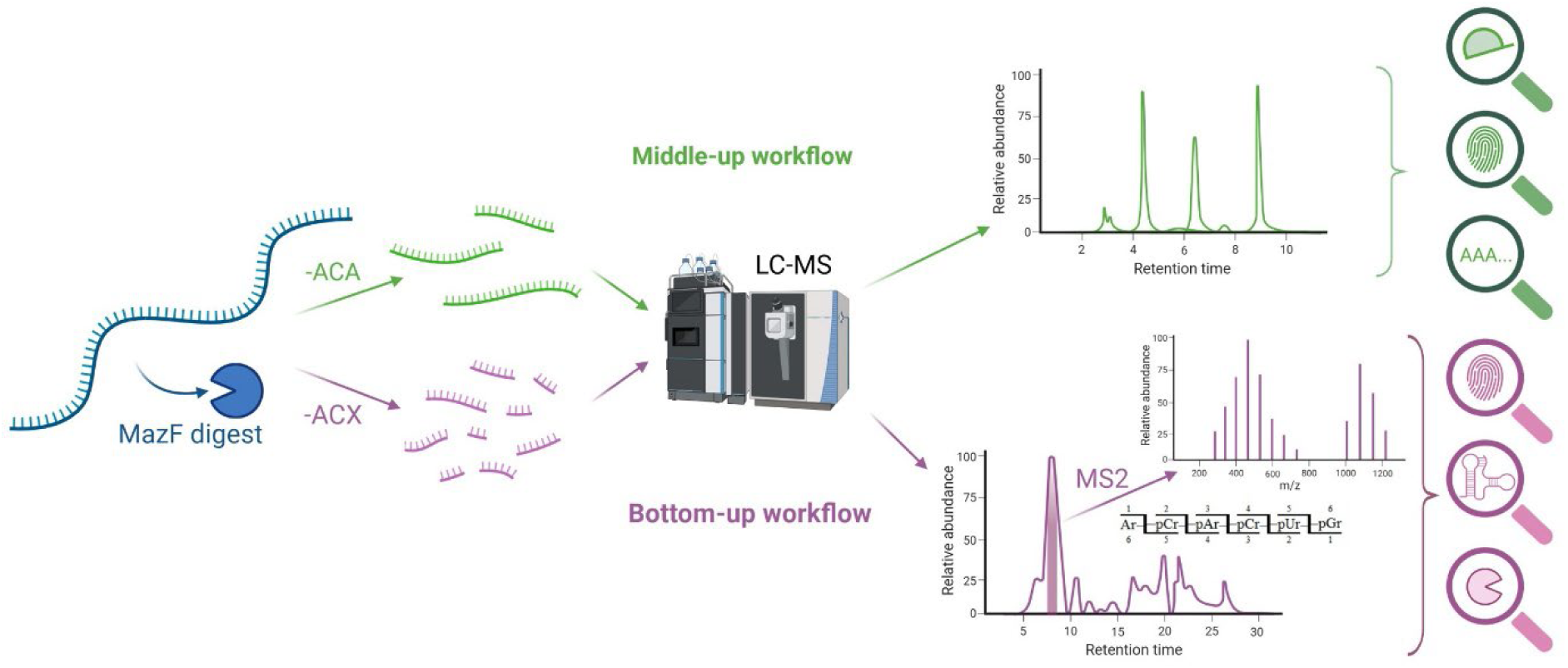
Schematic of middle-up versus bottom-up mRNA analysis workflow. In the middle-up workflow (green) *E. coli* MazF is optimised to cleave at the canonical recognitions sequence (-ACA). LC-MS analysis is then employed for intact mass analysis (MS1) enabling oligoribonucleotide mass fingerprinting (mRNA identity), 5’ capping efficiency and poly(A) tail characterisation. In the bottom-up workflow (purple) MazF digestions are optimised to cleave outside of the canonical recognition sequence resulting in (-ACX) cleavage. LC-MS/MS is then employed for comprehensive mRNA sequence mapping.

### Multi-attribute characterisation of mRNA using a middle-up LC-MS workflow

The middle-up workflow employs *E. coli* MazF, with mRNA digestion directed towards its canonical recognition sequence 5’-ACA via optimisation of the enzymatic digest conditions. Optimisation of the mRNA:MazF ratio, digest buffer and time of the reaction was performed to generate a series of larger oligoribonucleotide fragments at the canonical recognition sequence 5’-ACA. The confined fragment set allows for the development of a rapid QC method, with the oligoribonucleotides separated on an LC gradient in under 10 minutes.

Following rapid separation of the oligoribonucleotide fragments, we perform RNase mass mapping. By comparing these to a data set of molecular weights of oligoribonucleotides, produced by a theoretical 5’-ACA digest of our mRNA of interest, we can confirm the identity of the mRNA by matching the measured intact masses with the theoretical masses.

To facilitate accurate mass analysis of the high molecular weight oligonucleotides produced by 5’-ACA digests, it was necessary to optimise the parameters of the mass spectrometer. The Orbitrap Exploris 240 mass spectrometer was operated in the Intact Protein application mode with Low Pressure settings. Additionally, the Orbitrap resolution was set to 15,000. The combination of these parameters was used to minimise the loss of transient signal associated with high molecular weight analytes and enables the identification of RNA fragments up to approximately 200-300 nts in length by average mass.

The application of the middle-up LC-MS workflow for the rapid, multi-attribute characterisation of NLuc mRNA (∼871 nt) is shown in Figure 2. The corresponding LC-UV chromatogram generated via MazF digestion of NLuc mRNA optimised for cleavage at 5’-ACA of NLuc mRNA (∼871 nt) is shown in Figure 2A and the identified oligoribonucleotide fragments based on intact mass analysis are highlighted. The mass spectra and deconvoluted masses of two of the identified oligoribonucleotide fragments (267–315) and (316–416) are shown in Figures 2B/C respectively. A summary of all identified oligoribonucleotide fragments is shown in Table 1. The results show the identification of 12 different oligoribonucleotide fragments containing zero (7) or one (5) missed cleavage, resulting in 100% sequence coverage of NLuc mRNA based on oligoribonucleotide mass mapping.

**Figure 2:**
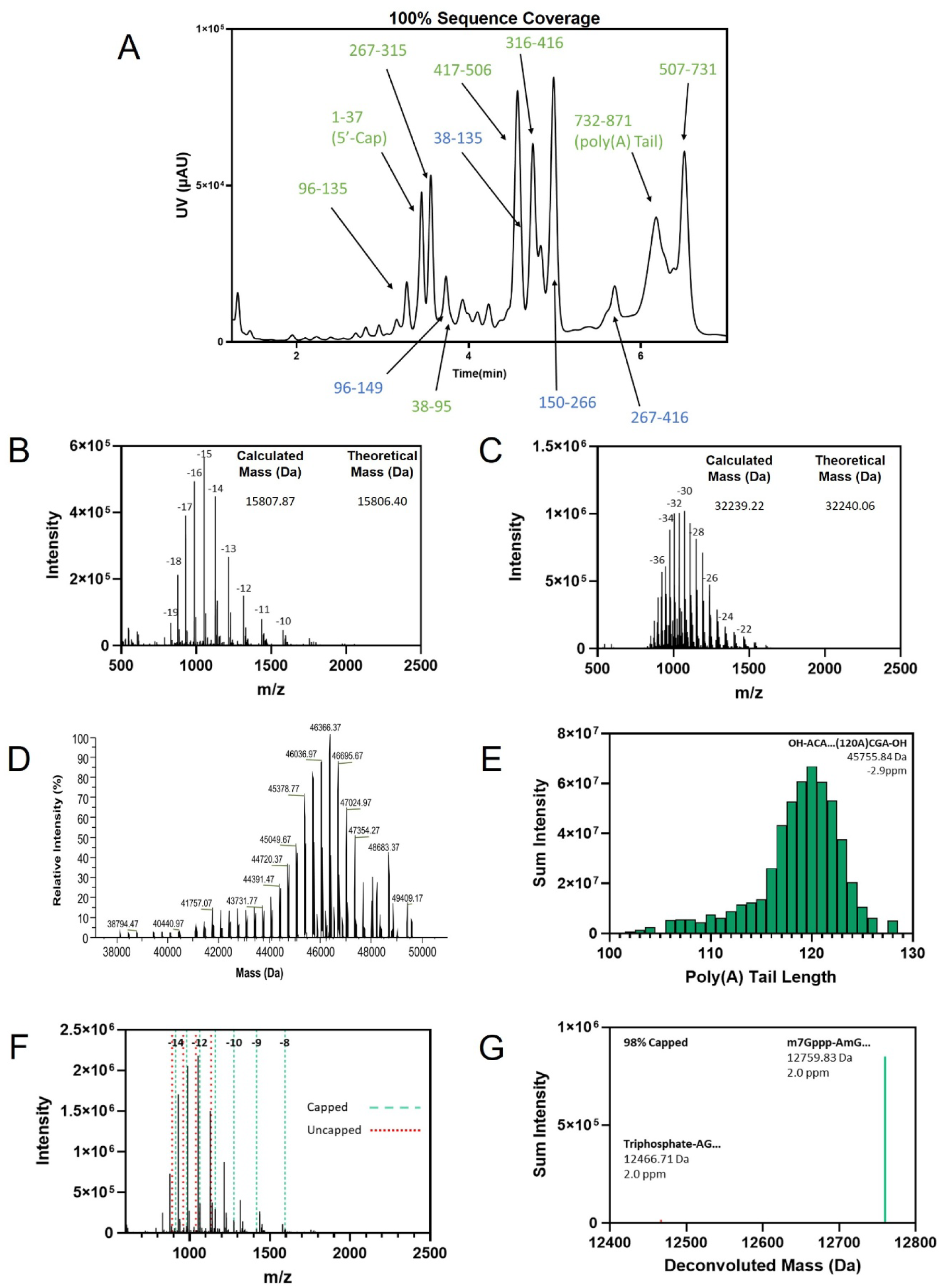
Middle-up MazF analysis of NLuc mRNA. (A) LC-UV chromatogram of a MazF digestion of NLuc mRNA. Oligoribonucleotide fragments identified from the intact mass analysis with 0 (green) and 1 (blue) missed cleavages are highlighted. (B) Example average spectrum from the chromatographic peak containing the 267-315 MazF fragment, annotated with the calculated average mass and theoretical mass. (C) Example average spectrum from the chromatographic peak containing the 316-416 MazF fragment, annotated with the calculated average mass and theoretical mass. (D) Deconvoluted average spectrum across the chromatographic peak containing the 3’-terminal MazF digest fragments. (E) Identified poly(A) containing 3’-terminal MazF digest fragments from LC-MS analysis. (F) Average spectrum of the chromatographic peaks containing the 5’-terminal MazF digest fragments. The charge states detected for the capped and uncapped species are shown in green and red respectively. (G) Deconvoluted spectrum of the identified 5’-terminal MazF fragments containing either the capped (cap-1) or the uncapped (triphosphate) end chemistries.

**Table 1-.** Identified MazF digest fragments generated from analysis of 5’-capped NLuc mRNA. Fragments in green and blue contain zero and one missed cleavage respectively.

| Fragment | Average Mass<br>(Da) | Expected Mass<br>(Da) | Mass Difference<br>(Da) |
| --- | --- | --- | --- |
| 1-37 | 12766.47 | 12765.71 | 0.76 |
| 96-135 | 12914.86 | 12914.63 | 0.23 |
| 267-315 | 15807.87 | 15806.40 | 1.47 |
| 96-149 | 17429.33 | 17430.33 | -1.00 |
| 38-95 | 18485.67 | 18486.88 | -1.20 |
| 417-506 | 29050.32 | 29051.28 | -0.95 |
| 38-135 | 31400.26 | 31401.51 | -1.25 |
| 316-416 | 32239.22 | 32240.06 | -0.84 |
| 150-266 | 37845.00 | 37845.58 | -0.58 |
| 732-871 | 45756.02 | 45755.54 | 0.48 |
| 267-416 | 48047.68 | 48046.46 | 1.21 |
| 507-731 | 72273.62 | 72271.73 | 1.89 |

The same LC-MS data set produced for the identity testing can also be used for additional mRNA characterisation, including analysis of the 5’ capping efficiency and 3’ poly(A) tail length and heterogeneity. By analysing the 3’ terminal oligoribonucleotide fragment (732– 871) produced by the MazF digest of the NLuc mRNA, we can determine the length and heterogeneity of the poly(A) tail. Figure 2D shows the deconvoluted mass spectrum produced by this oligoribonucleotide fragment; the MS spectrum of the poly(A) containing fragment is complex and includes a relatively wide range of masses. Deconvolution of this spectrum reveals a major profile of peaks, each of which are separated by the difference of an “A”, and the masses of each peak can be assigned to a set of different poly(A) lengths. In Figure 2E, we have plotted the identified lengths against the sum intensities of the corresponding MS peaks to generate a profile of the poly(A) tail heterogeneity. Thus, for this NLuc mRNA, a mean poly(A) length of 120 nt (the length prescribed by the DNA template) with a bivariate Gaussian distribution of lengths observed.

Figure 2F shows the MS spectra produced by the 5’ terminal oligoribonucleotide (1–37 fragment) produced by the MazF digest of NLuc mRNA. The results show more than one distinct charge state profile, indicating the presence of multiple species. Figure 2G shows the deconvoluted mass spectrum of this oligo, which yields two peaks of interest: one corresponding to the 5’ Cap-1 species (m7GpppAmG) and the other corresponding to an uncapped (pppAG) oligoribonucleotide fragment (1–37). Using the sum intensities of these two peaks, we determined a 5’ capping efficiency of 98% (see Figure S1).

To confirm that this method is also applicable to modified mRNAs, we analysed NLuc mRNA with uridine replaced with N1-methylpseudouridine via the middle-up LC-MS workflow. Here, we achieved 98% sequence coverage via identification of 12 oligoribonucleotide fragments (see Supporting Figure S2 and Table S2).

The same middle-up workflow was applied to eGFP mRNA for sequence identity and poly(A) tail analysis (see Supporting Figure S3). The LC-UV chromatogram generated from the 5’-ACA MazF digest, has a distinctly different chromatographic profile compared to the NLuc mRNA (Figure 2A), due to different oligoribonucleotide fragments generated.

Following deconvolution, analysis of the intact mass data generated by the workflow enabled the identification of the majority of oligoribonucleotide fragments from the LC-UV chromatogram with zero or one missed cleavage (see Table S2). The 16 identified of oligoribonucleotide resulted in a sequence coverage of 87% of the eGFP. The missing portion of this sequence comes from the 5’ terminal 1-123 oligonucleotide.

One of the identified oligoribonucleotide fragments corresponds to the 3’ terminal, poly(A) containing fragment (760–930 nt). Following deconvolution length and heterogeneity was determined for the eGFP mRNA (see Supporting Figure S3E).

Finally, we demonstrate the middle-up workflow on a longer mRNA: FLuc mRNA (∼1928 nt). Consistent with previous mRNA analysis, the LC-UV profile shows a chromatographic profile that is distinct from the previous LC UV chromatograms generated from MazF digestion of NLuc and eGFP mRNAs. Data analysis led to the identification of 40 different oligoribonucleotide fragments containing zero or one missed cleavage (Table 2), which culminate in a total FLuc mRNA sequence coverage of 74%. The average mass spectra of identified oligoribonucleotide fragments from peak X (548–643) and peak Y (755–958) are shown in Figures 3B&C respectively. The MS spectra for peak Y shows multiple overlapping charge series and deconvolutes to multiple masses of predicted 5’-ACA digest fragments (42777.18 Da, 43280.59 Da, 44322.83 Da, 44872.53 Da), demonstrating the co-elution and identification of the 4 oligoribonucleotide fragments (380-511, 620-754, 755-892, and 1124-1261) in this peak. The missing sequence from this digest corresponds to the 1442-1928 fragment at the 3’ terminus. This region is deficient in 5’-ACA motifs, resulting in a large oligonucleotide fragment not amenable to MS analysis with this workflow. As such, it was not possible to assess the length and heterogeneity of the poly(A) tail from this digest.

**Figure 3:**
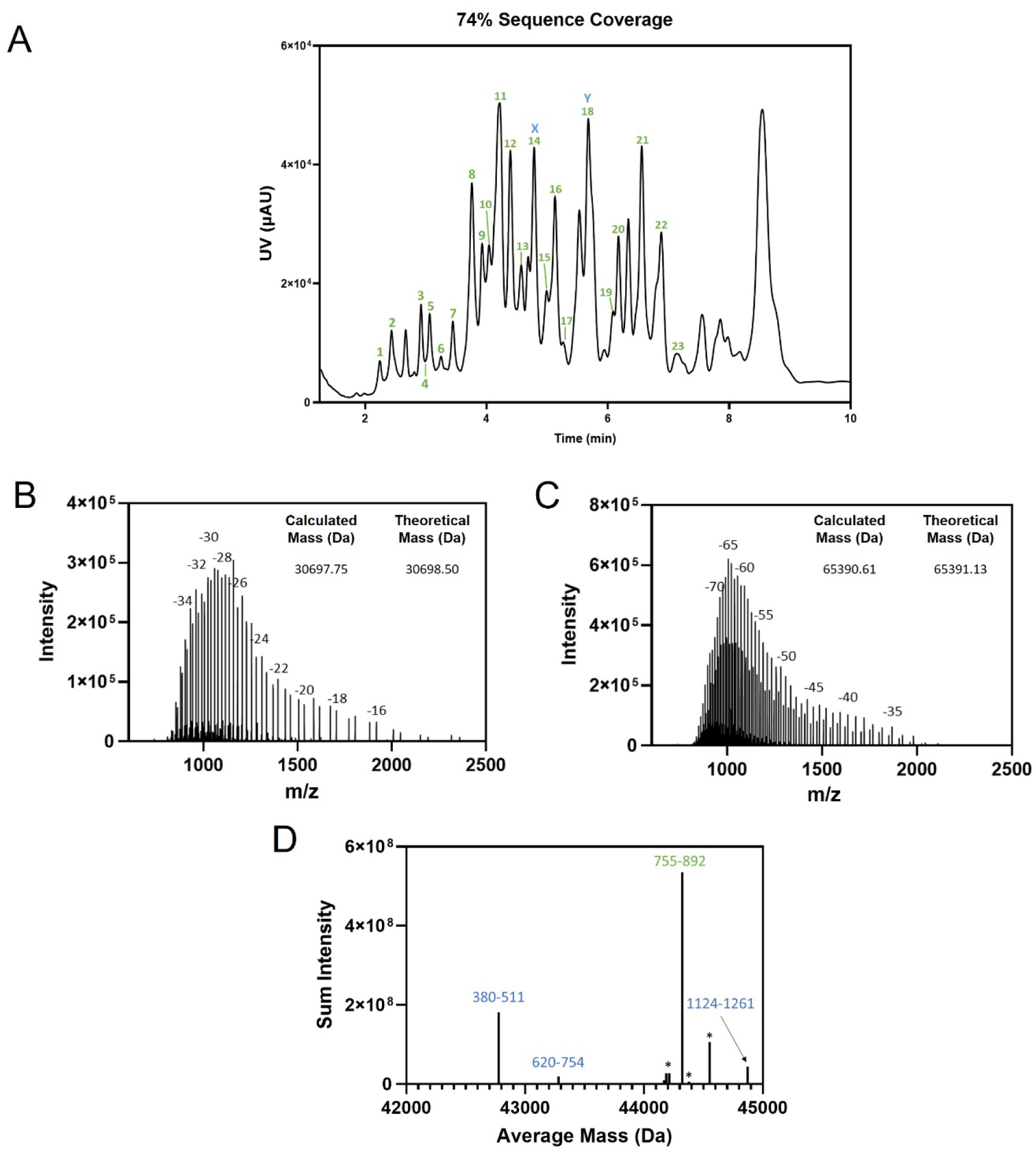
Middle-up MazF analysis of an FLuc mRNA. (A) LC-UV chromatogram of an *E. coli* MazF digestion of NLuc mRNA annotated with identified 0 and 1 missed cleavage species identified from LC-MS analysis, numbered according to table 2. (B) Example average spectrum from the chromatographic peak containing the 548-643 MazF fragment (peak X), annotated with the calculated average mass and theoretical mass. (C) Example average spectrum from the chromatographic peak containing the 755-958 MazF fragment (peak Y), annotated with the calculated average mass and theoretical mass. (D) Deconvoluted average mass spectrum from peak Y, annotated with identified 0 (green) and 1 (blue) missed cleavges. Unidentified masses hypothesised to correspond to >1 missed cleavage and non-ACA cleavage are also annotated (*)

**Table 2-.** Identified MazF digest fragments from an FLuc mRNA. Fragments in green and blue contain zero and one missed cleavage respectively.

| Peak | Fragment(s) | Average Mass<br>(Da) | Expected Mass<br>(Da) | Mass Difference<br>(Da) |
| --- | --- | --- | --- | --- |
| 1 | 620-643 | 7680.80 | 7680.656 | 0.15 |
| 2 | 1292-1318 | 8824.62 | 8826.363 | -1.74 |
| 3 | 1262-1291 | 9589.84 | 9589.84 | 0.00 |
| 4 | 512-547 | 11615.86 | 11616.049 | -0.19 |
| 5 | 272-304 | 10691.73 | 10691.487 | 0.25 |
| 6 | 30-70 | 13313.66 | 13315.164 | -1.51 |
| 7 | 1262-1309 | 15461.43 | 15461.397 | 0.03 |
| 8 | 134-190 | 18266.64 | 18267.033 | -0.39 |
|  | 1385-1441 | 18371.61 | 18370.123 | 1.49 |
| 9 | 1-29 | 9492.29 | 9491.461 | 0.83 |
|  | 1124-1183 | 19499.36 | 19497.715 | 1.65 |
|  | 71-133 | 20242.81 | 20241.252 | 1.55 |
| 10 | 893-958 | 21069.17 | 21067.648 | 1.52 |
|  | 1319-1384 | 21298.13 | 21299.824 | -1.69 |
| 11 | 305-373 | 22081.84 | 22080.186 | 1.66 |
|  | 134-202 | 22203.69 | 22203.41 | 0.28 |
|  | 203-271 | 22351.21 | 22349.481 | 1.72 |
|  | 548-619 | 23019.43 | 23017.843 | 1.59 |
| 12 | 305-379 | 23962.89 | 23961.31 | 1.58 |
|  | 1310-1384 | 24255.37 | 24254.63 | 0.74 |
|  | 1184-1261 | 25373.63 | 25375.327 | -1.70 |
| 13 | 1-70 | 22808.14 | 22806.625 | 1.51 |
|  | 191-271 | 26284.12 | 26285.859 | -1.74 |
| 14 (X) | 548-643 | 30697.75 | 30698.499 | -0.75 |
| 15 | 272-373 | 32772.08 | 32771.673 | 0.41 |
| 16 | 512-619 | 34633.18 | 34633.892 | -0.71 |
|  | 1184-1291 | 34965.22 | 34965.167 | 0.05 |
|  | 644-754 | 35599.72 | 35600.494 | -0.77 |
| 17 | 401-511 | 35901.79 | 35902.729 | -0.94 |
|  | 71-190 | 38507.61 | 38508.285 | -0.68 |
| 18 (Y) | 380-511 | 42777.18 | 42777.913 | -0.73 |
|  | 620-754 | 43280.59 | 43281.15 | -0.56 |
|  | 755-892 | 44322.83 | 44323.477 | -0.65 |
|  | 1124-1261 | 44872.53 | 44873.042 | -0.51 |
| 19 | 401-547 | 47518.73 | 47518.778 | -0.05 |
| 20 | 959-1123 | 53324.66 | 53325.295 | -0.64 |
| 21 | 755-958 | 65390.61 | 65391.126 | -0.52 |
| 22 | 959-1183 | 72822.59 | 72823.01 | -0.42 |
|  | 893-1123 | 74392.70 | 74392.944 | -0.24 |
| 23 | 644-892 | 79923.95 | 79923.971 | -0.02 |

### Characterisation of mRNA using MazF digestions in conjunction with a bottom-up LC-MS workflow

In the bottom-up workflow, we aimed to generate a larger number of oligoribonucleotide fragments with a smaller size distribution by exploiting less specific MazF digestion (5’-ACX). This was accomplished by re-optimising the digest buffer optimisation and time of the reaction to generate larger numbers of smaller sized oligoribonucleotide fragments. Following MazF digestion, the oligoribonucleotides were separated using IP-RP HPLC interfaced with MS/MS analysis.

The LC-UV chromatogram of the MazF digest of NLuc mRNA under the optimised 5’-ACX digestions conditions is shown in Figure 4A. The results show the generation of a wide range of smaller oligoribonucleotide fragments (<50 mers) compared to the previous MazF digest (see Figure 2A and Figure 4A).

**Figure 4:**
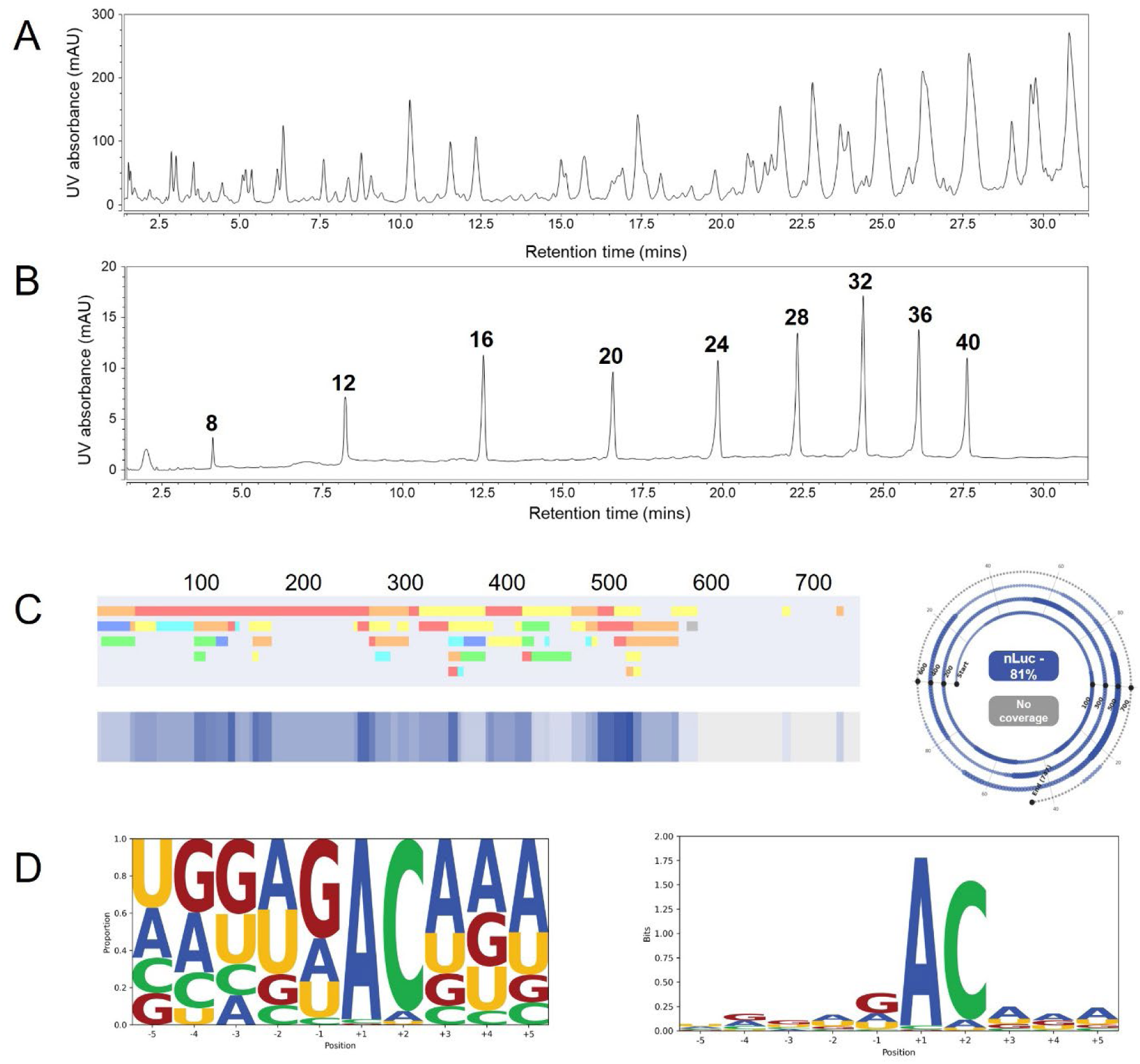
Bottom-up MazF analysis of Nluc mRNA. (A) LC-UV chromatogram of an -ACX *E. coli* MazF digestion of NLuc mRNA. (B) LC-UV chromatogram of an 8-40-mer standard. (C) Linear and spiral sequence maps generated for the NLuc mRNA digest. (D) Sequence logos representing the cleavage specify of MazF under –ACX digest conditions, where the cleavage site is between the −1 and +1 base positions. RNA sequences flanking inferred cleavage sites were aligned and nucleotide frequencies at positions −5 to −1 and +1 to +5 were calculated. Logos display either nucleotide proportions (left) or information content (right), calculated from position-specific probabilities relative to a uniform background.

Oligoribonucleotide identifications were performed using automated data analysis software to identify oligoribonucleotides on the basis of their accurate mass in conjunction with the MS/MS fragmentation spectra and map the corresponding oligoribonucleotide sequences to the known RNA sequence.^15,19^ mRNA sequence coverage was determined using only unique oligoribonucleotide fragment identifications (see Tables S1 and S4) and the percentage coverage was determined from the mRNA sequence (not including the poly(A) tail). Furthermore, high sequence coverages with no or low sequence matches against random control RNA sequences were obtained, demonstrating the specificity of the analytical workflow in conjunction with the parameters used for mRNA sequence mapping (see Table S4).

The LC-MS/MS analysis of the MazF digest of NLuc mRNA resulted in 81% sequence coverage (based only on unique oligonucleotide identifications) using the optimised 5’-ACX digestion conditions, the corresponding mRNA sequence map is shown in Figure 4C. Further analysis of the mRNA sequence coverage and identified oligoribonucleotide fragments reveals that unmapped sections of the mRNA sequence contain limited incidences of 5’-ACX motifs, leading to theoretical fragments >50 nt in length. The fragmentation and subsequent identification of oligoribonucleotide sequences is generally limited to oligoribonucleotides of <50 nt.

Further examination of the LC MS/MS data was performed in BioPharmaFinder by comparing the identified oligoribonucleotides from nonspecific cleavage search of NLuc mRNA. The results provide further insight int the cleavage specificity of MazF under the optimised 5’-ACX digest conditions. The sequence logos in Figure 4D demonstrate that, as expected, the preferential cleavage occurs at 5’-AC, with only a small preference for cleavage at 5’-ACA. Interestingly, the logos also reveal that there are limited incidences of cleavage 3’ of a C and there is a slight preference for cleaving 3’ of a G. This is not representative of the number of theoretical 5’-ACX cleavage sites for NLuc: 10 5’-ACA sites, 10 5’-ACC sites, 12 5’-ACG sites and 24 5’-ACU sites. The list of oligoribonucleotide dentifications annotated with bases either side of the cleavage position can be found in Supporting Table S1 and a position frequency and probability matrices for the data set can be found in Supporting Table S5 and S6.

### LC-MS/MS sequence mapping of CSP mRNA

The bottom-up workflow for direct mRNA sequence mapping analysis was also performed on SARS-CoV-2 Spike Protein (CSP) mRNA (∼4300 nt). Obtaining high sequence coverage based on unique oligoribonucleotide fragments is more challenging for larger mRNA sequences and those mRNAs with high degree of secondary structure. Improved sequence coverage of larger mRNA constructs that have more significant secondary structure, such as CSP mRNA (>4000 nt), can be achieved by implementing multiple RNases and combining LC-MS/MS data sets.^14,19^ Therefore, in this work we utilise LC-MS/MS data generated by complementary enzymes RNase T1 and RNase U2 using partial RNase digests,^19^ in combination with MazF digest data. The corresponding LC-UV chromatograms for each mRNA digest are shown in Figure 5A, highlighting the differences in oligoribonucleotide fragments generated via the different enzymes.

**Figure 5:**
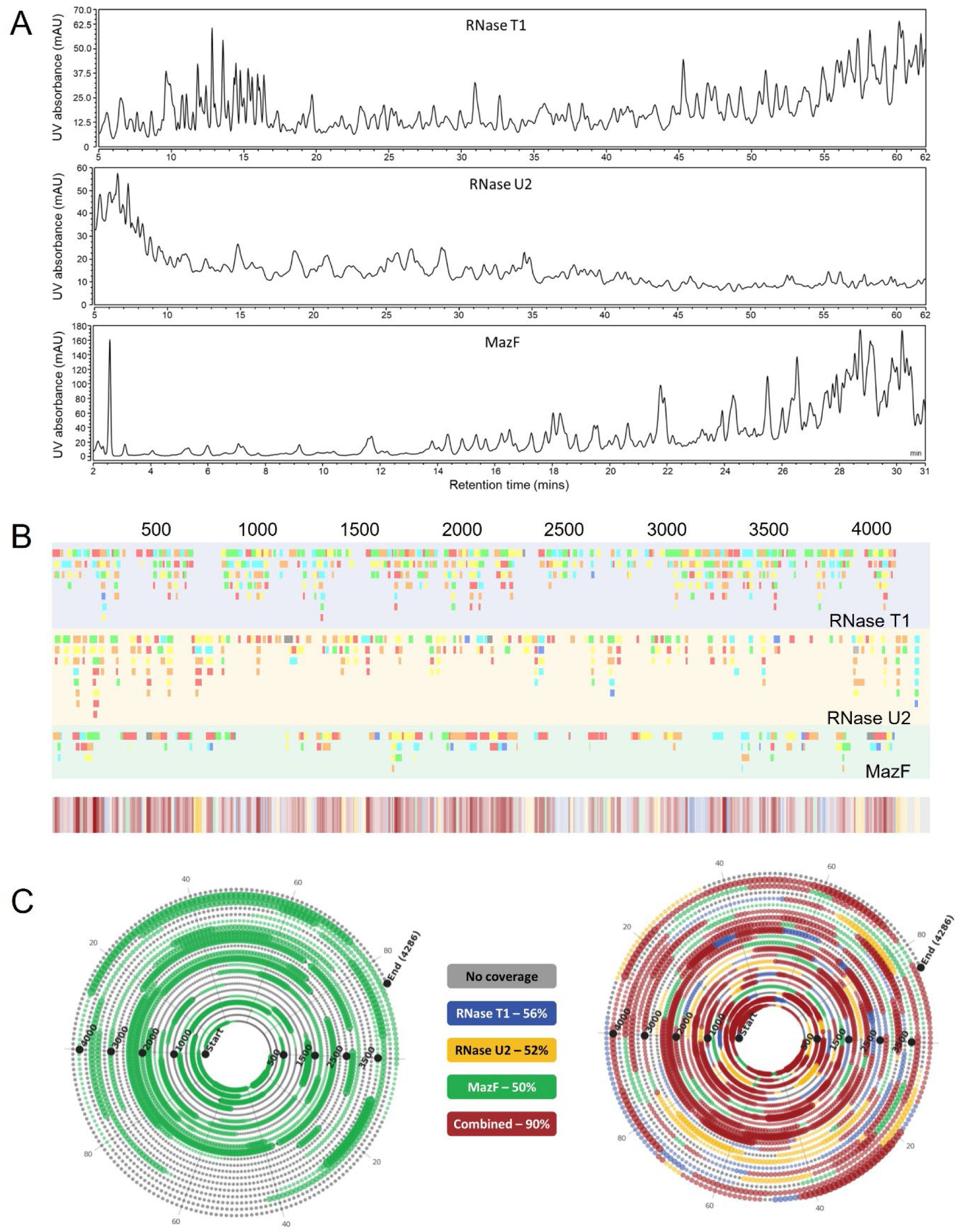
Bottom-up analysis of a CSP mRNA. (A) LC-UV chromatograms of an RNase T1 (top), an RNase U2 (middle) and a MazF digestion (bottom) of CSP mRNA. (B) Linear sequence maps produced by RNase T1 (blue), RNase U2 (yellow) and MazF (green). (C) Spiral plots of the sequence coverage identified for the MazF digest (left) and the combined complementary digests (right).

To better visualise the sequencing data, we implement our in-house sequence mapping software tools.^19^ The results of the differences in unique oligoribonucleotide identifications are further evident in the linear sequences maps (Figure 5B), where a coloured bar represents each oligo fragment identification that is maps onto the mRNA sequence. This map reveals regions of the CSP mRNA that are universally mapped by all three enzymes, and areas where complementary digests provide overlapping coverage to fill gaps. The benefit of using multiple enzymes is further evident when we convert the oligo mapping into mRNA sequence coverage. In the left-hand spiral plot shown in Figure 5C, the mRNA sequence coverage that was obtained using MazF enzyme alone a sequence coverage of 50% was obtained via the bottom-up MazF workflow. Moreover, by employing all three complementary enzymes, the mRNA sequence coverage of CSP mRNA was increased to 90% (right-hand plot in Figure 5C).

## Conclusions

We have developed two complementary LC-MS-based workflows exploiting MazF’s site-specific cleavage for comprehensive mRNA characterization. By manipulating the reaction conditions, we have optimised for cleavage at *E. coli* MazF’s canonical recognition sequence, 5’-ACA, for rapid mRNA mass mapping in conjunction with intact mass analysis, and redirected to less specific, 5’-ACX cleavage for detailed mRNA sequence mapping via MS/MS.

The middle-up workflow addresses a critical need for rapid, multi-attribute QC that is suitable for manufacturing environments. Achieving complete assessment—sequence identity, 5’ capping efficiency, and poly(A) tail analysis—in under 10 minutes represents a significant efficiency gain over traditional approaches. The successful application to NLuc (100% coverage), eGFP (87% coverage), and modified NLuc mRNA (98% coverage) demonstrates versatility across different constructs and modifications. Incomplete FLuc coverage (74%) highlights a sequence-dependent limitation: regions deficient in “-ACA” motifs generate fragments too large for conventional MS analysis under the conditions used in this study. Further optimisation of MS conditions may enable detection of higher molecular weight fragments and improve coverage. *In silico* analysis could be implemented to identify such regions and inform method selection. However, coverage here remains sufficient to confirm construct identity through detection of unique sequence fragments, namely across the coding region of the transcript. The ability to simultaneously assess multiple CQAs from a single analysis represents significant efficiency gains. Our determination of 5’ capping efficiency (98% for NLuc) and comprehensive poly(A) tail profiling directly from MazF digests eliminates the need for separate assays.

Our bottom-up workflow demonstrates that *E. coli* MazF can be used for mRNA sequence mapping under modified reaction conditions to less specific, 5’-ACX cleavage, generating smaller oligoribonucleotide fragments. The 50% coverage of CSP mRNA via MazF alone, extended to 90% when combined with RNase T1 and U2, illustrates the power of complementary digests. Minimal false-positive identifications against random sequences confirm the robustness of our identification criteria.

These workflows are readily adaptable to emerging modifications, as demonstrated with N1-methylpseudouridine-modified mRNA. As the field evolves toward increasingly complex constructs, the flexibility to switch between rapid QC and detailed characterisation modes will become invaluable.

In conclusion, MazF-based MS workflows provide versatile, orthogonal tools bridging high-throughput analysis requirements and comprehensive characterisation studies, representing a significant addition to the analytical toolkit supporting mRNA-based vaccine and therapeutic development.

## Supporting Information

- Supporting Figure S1. Extracted ion chromatograms of NLuc 5’-terminal fragments
- Supporting Figure S2. Middle-up MazF analysis of a capped and N1-methylpseudouridine modified NLuc mRNA
- Supporting Figure S3 Middle-up MazF analysis of an eGFP mRNA
- Supporting Table S1 Identified oligoribonucleotides using LC MS/MS.
- Supporting Table S2. Identified MazF digest fragments from a capped and chemically modified NLuc mRNA
- Supporting Table S3. Identified MazF digest fragments from eGFP mRNA
- Supporting Table S4. mRNA sequence coverage using mRNA sequence mapping with MazF
- Supporting Table S5. Position frequency matrix for the cleavage sites in a MazF digest of NLuc mRNA
- Supporting Table S6: Position probability matrix for the cleavage sites in a MazF digest of NLuc mRNA

## Acknowledgements

MJD and ZK acknowledge funding from the Wellcome Leap R3 programme; Coalition for Epidemic Preparedness Innovations (CEPI) and UK Research and Innovation/ Biotechnology and Biological Sciences Research Council (BBSRC), Engineering Biology Mission Award [BB/Y007514/1]. GRO is funded through a University of Sheffield Engineering and Physical Sciences Research Council (EPSRC) Doctoral Training Partnership Institute for Sustainable Food Scholarship [EP/T517835/1]. LPW is funded through a BBSRC White Rose DTP CASE Studentship in collaboration AstraZeneca.

