## Supporting Information for "Multi-attribute characterisation of mRNA via MazF endoribonuclease and LC-MS workflows"

### Table of Contents:

Supporting Figure S1. Extracted ion chromatograms of NLuc 5'-terminal fragments

Supporting Figure S2. Middle-up MazF analysis of a capped and N1-methylpseudouridine modified NLuc mRNA

Supporting Figure S3 Middle-up MazF analysis of an eGFP mRNA

Supporting Table S2. Identified MazF digest fragments from a capped and chemically modified NLuc mRNA

Supporting Table S3. Identified MazF digest fragments from eGFP mRNA

Supporting Table S4. mRNA sequence coverage using mRNA sequence mapping with MazF

Supporting Table S5. Position frequency matrix for the cleavage sites in a MazF digest of NLuc mRNA

Supporting Table S6: Position probability matrix for the cleavage sites in a MazF digest of NLuc mRNA

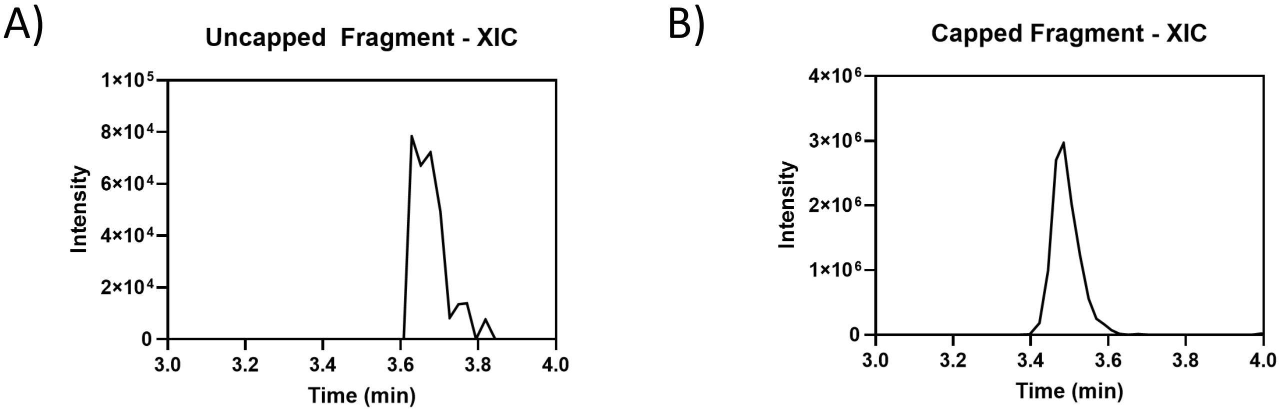


Supporting Figure S1: Extracted ion chromatograms of NLuc 5′-terminal fragments. (A) Extracted ion chromatogram of the $12^-$ charge state of the uncapped (triphosphate) 1-37 MazF digest fragment ±10 ppm. (C) Extracted ion chromatogram of the $10^-$ charge state of the capped (cap-1) 1-37 MazF digest fragment ±10 ppm.

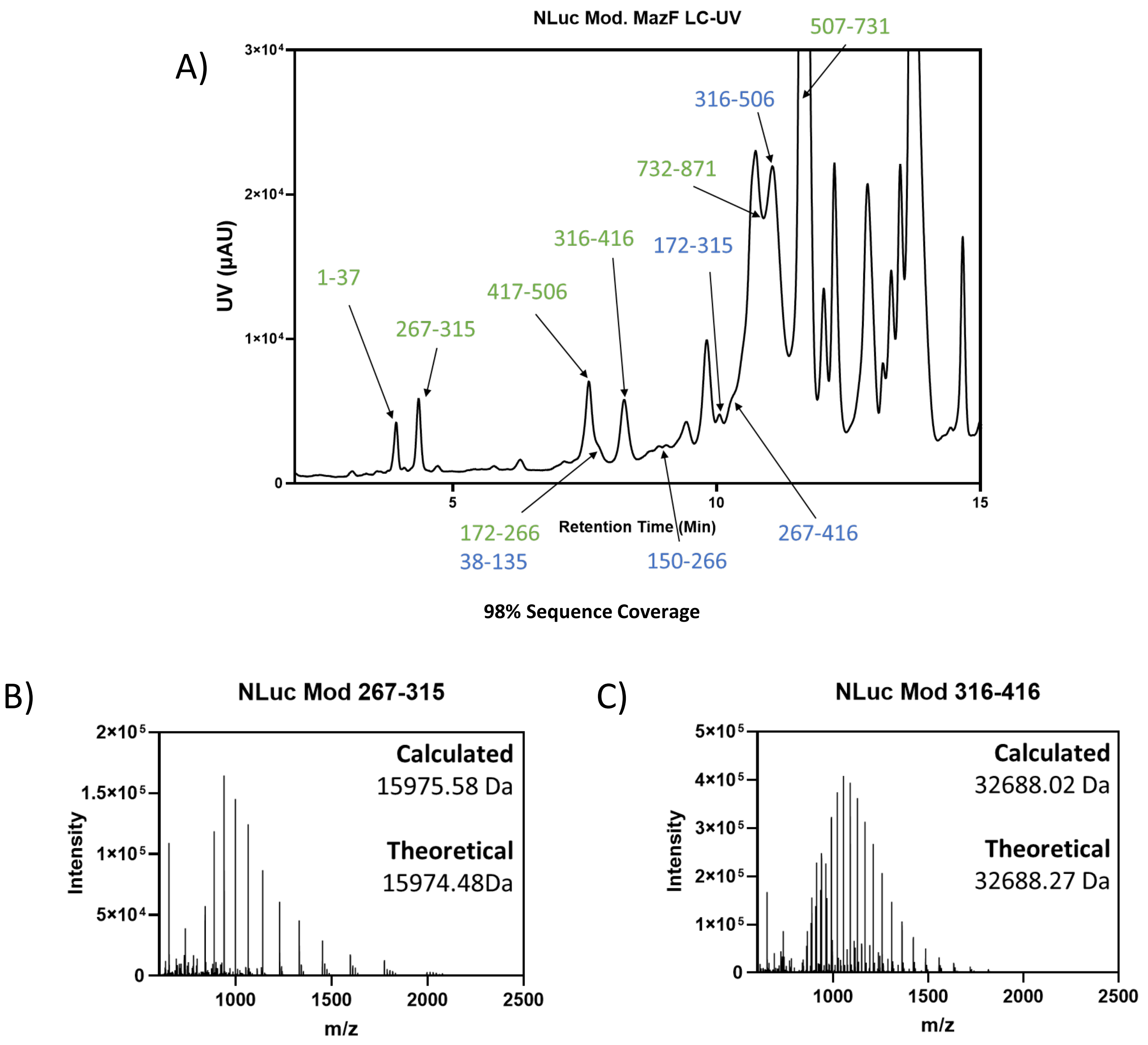


| Calculated | Theoretical |
|---|---|
| 15975.58 Da | 15975.58 Da |

| Calculated | Theoretical |
|---|---|
| 32688.02 Da | 32688.27 Da |

Supporting Figure S2. Middle-up MazF analysis of a capped and N1-methylpseudouridine modified NLuc mRNA. (A) LC-UV chromatogram of a MazF digestion of NLuc mRNA annotated with identified 0 (green) and 1 (blue) missed cleavage species identified from LC-MS analysis. (B) Example average spectrum from the chromatographic peak containing the 267-315 MazF fragment, annotated with the calculated average mass and theoretical mass. (C) Example average spectrum from the chromatographic peak containing the 267-315 MazF fragment, annotated with the calculated average mass and theoretical mass.

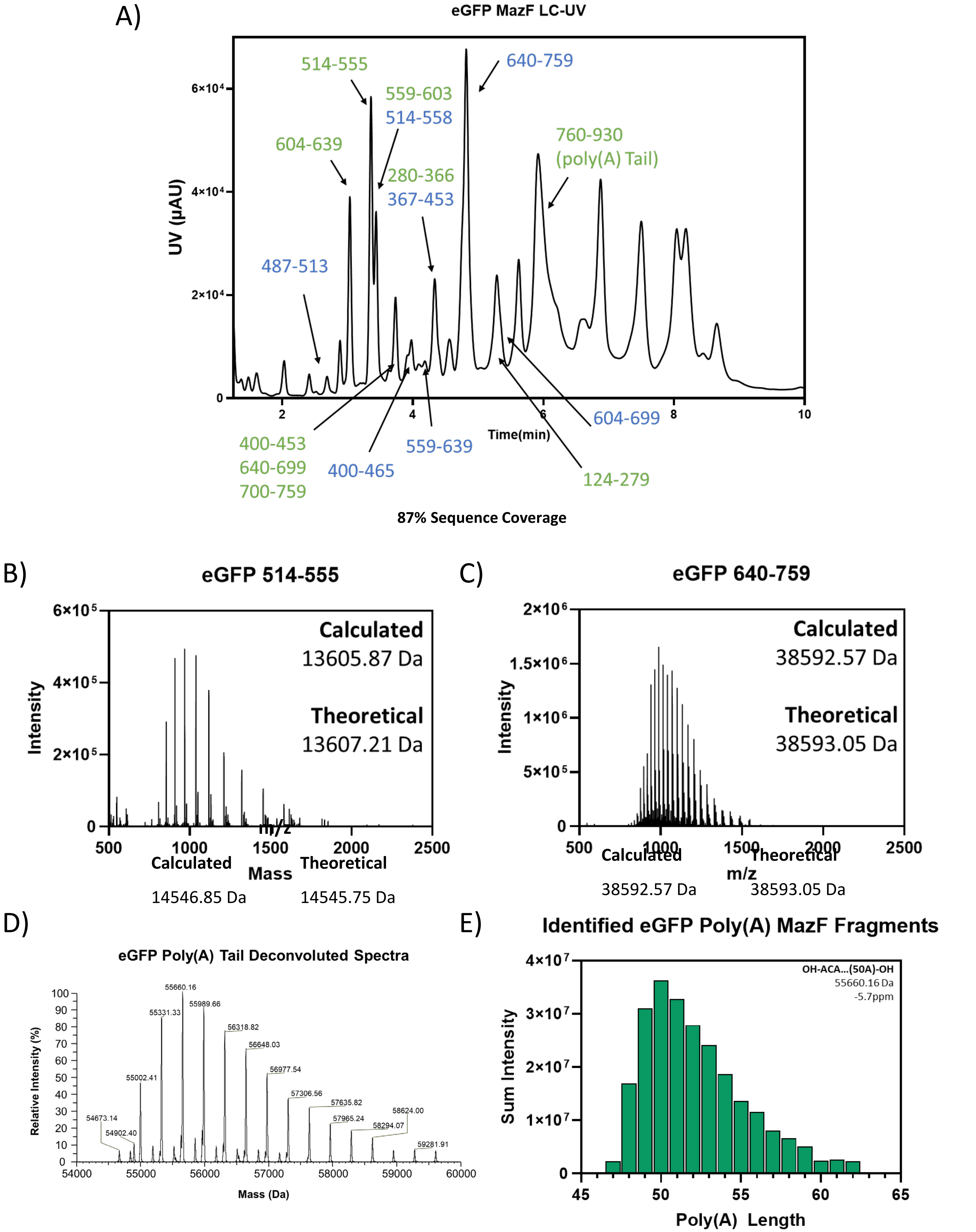


Supporting Figure S3. Middle-up MazF analysis of an eGFP mRNA. (A) LC-UV chromatogram of a MazF digestion of NLuc mRNA annotated with identified 0 (green) and 1 (blue) missed cleavage species identified from LC-MS analysis. (B) Example average spectrum from the chromatographic peak containing the 559-603 MazF fragment, annotated with the calculated average mass and theoretical mass. (C) Example average spectrum from the chromatographic peak containing the 640-759 MazF fragment, annotated with the calculated average mass and theoretical mass. (D) Deconvoluted average spectrum across the chromatographic peak containing the 3'-terminal MazF digest fragments. (E) Identified poly(A) containing 3'-terminal MazF digest fragments from LC-MS analysis.

Supporting Table S2. Identified MazF digest fragments from a 5' capped and chemically modified NLuc mRNA. Deconvoluted average masses of oligonucleotide digest fragments identified from a 6 hr digest, supplemented with 40U MazF after 3 hours, of NLuc mRNA co-transcriptionally capped with CleanCap™ AG capping reagent and transcribed with N1-methylpseudouridine. Oligonucleotides containing zero missed cleavages (at ^ACA) are shown in green, and oligonucleotides containing one missed cleavage are shown in blue.

| Fragment | Average Mass | Expected | Mass Difference (Da) | Error (ppm) |
|---|---|---|---|---|
| 1-37 | 12822.39 | 12821.74 | 0.66 | 51.38 |
| 267-315 | 15975.58 | 15974.48 | 1.10 | 68.95 |
| 417-506 | 29387.07 | 29387.44 | -0.36 | -12.39 |
| 172-266 | 31066.94 | 31067.49 | -0.54 | -17.48 |
| 38-135 | 31808.66 | 31807.70 | 0.97 | 30.36 |
| 316-416 | 32688.02 | 32688.27 | -0.25 | -7.75 |
| 150-266 | 38209.70 | 38209.75 | -0.05 | -1.43 |
| 732-871 | 45880.95 | 45881.60 | -0.65 | -14.17 |
| 172 -315 | 47041.74 | 47041.97 | -0.22 | -4.76 |
| 267-416 | 48662.67 | 48662.76 | -0.08 | -1.74 |
| 316-506 | 62076.08 | 62075.71 | 0.37 | 5.99 |
| 507-731 | 73224.61 | 73224.19 | 0.43 | 5.81 |

Supporting Table S3. Identified MazF digest fragments from eGFP mRNA. Deconvoluted average masses of oligonucleotide digest fragments identified from a 4 hr digest of eGFP mRNA. Oligonucleotides containing zero missed cleavages (at ^ACA) are shown in green, and oligonucleotides containing one missed cleavage are shown in blue.

| Fragment | Average Mass | Expected | Mass Difference (Da) | Error (ppm) |
|---|---|---|---|---|
| 487-513 | 8638.36 | 8637.16 | 1.20 | 138.37 |
| 604-639 | 11486.86 | 11486.84 | 0.02 | 1.70 |
| 514-555 | 13605.87 | 13607.21 | -1.34 | -98.48 |
| 559-603 | 14546.85 | 14545.75 | 1.10 | 75.56 |
| 514-558 | 14572.12 | 14570.81 | 1.31 | 89.91 |
| 400-453 | 17528.76 | 17527.51 | 1.25 | 71.43 |
| 640-699 | 19254.24 | 19252.61 | 1.62 | 84.39 |
| 700-759 | 19341.94 | 19340.43 | 1.51 | 78.00 |
| 400-465 | 21401.43 | 21399.81 | 1.61 | 75.45 |
| 559-639 | 26031.99 | 26032.58 | -0.60 | -22.90 |
| 280-366 | 27920.14 | 27918.67 | 1.47 | 52.68 |
| 367-453 | 28312.63 | 28313.99 | -1.36 | -48.08 |
| 604-699 | 30737.73 | 30739.45 | -1.72 | -55.80 |
| 640-759 | 38592.57 | 38593.05 | -0.48 | -12.42 |
| 124-279 | 50025.95 | 50025.78 | 0.17 | 3.50 |
| 760-930 | 55002.50 | 55002.06 | 0.44 | 7.99 |

Supporting Table S4. mRNA sequence coverage using mRNA sequence mapping. Sequence coverages are shown for different mRNAs and randomised RNA of the same length and GC content for the LC-MS/MS analysed enzymatic digests of NLuc and CSP mRNA.

| mRNA | Nuclease | Sequence coverage / % | | |
|---|---|---|---|---|
| | | mRNA (unique only) | mRNA (inc. non-uniques) | Randomised (unique only) |
| NLuc | MazF | 80.6 | 80.6 | 0.0 |
| CSP | MazF | 50.4 | 50.8 | 0.0 |
| | RNase T1 | 55.8 | 55.9 | 0.0 |
| | RNase U2 | 52.5 | 53.4 | 1.1 |

Supporting Table S5. Position frequency matrix for the cleavage sites in a MazF digest of NLuc mRNA. Cleavage is between the -1 and +1 base positions

| | -5 | -4 | -3 | -2 | -1 | 1 | 2 | 3 | 4 | 5 |
|---|---|---|---|---|---|---|---|---|---|---|
| A | 38 | 51 | 35 | 45 | 57 | 143 | 32 | 82 | 64 | 66 |
| C | 31 | 27 | 30 | 16 | 10 | 9 | 112 | 16 | 22 | 26 |
| G | 36 | 47 | 45 | 35 | 57 | 3 | 3 | 29 | 32 | 26 |
| U | 47 | 28 | 48 | 59 | 31 | 0 | 8 | 28 | 37 | 37 |

Supporting Table S6. Position probability matrix for the cleavage sites in a MazF digest of NLuc mRNA. Cleavage is between the -1 and +1 base positions

| | -5 | -4 | -3 | -2 | -1 | 1 | 2 | 3 | 4 | 5 |
|---|---|---|---|---|---|---|---|---|---|---|
| A | 0.27 | 0.32 | 0.16 | 0.38 | 0.23 | 0.97 | 0.05 | 0.51 | 0.39 | 0.50 |
| C | 0.19 | 0.19 | 0.17 | 0.11 | 0.04 | 0.02 | 0.93 | 0.10 | 0.08 | 0.12 |
| G | 0.17 | 0.40 | 0.40 | 0.17 | 0.54 | 0.01 | 0.00 | 0.17 | 0.29 | 0.15 |
| U | 0.37 | 0.09 | 0.27 | 0.35 | 0.19 | 0.00 | 0.02 | 0.22 | 0.24 | 0.23 |
